# High-resolution structural genomics reveals new therapeutic vulnerabilities in glioblastoma

**DOI:** 10.1101/442277

**Authors:** Michael J Johnston, Ana Nikolic, Nicoletta Ninkovic, Paul Guilhamon, Florence MG Cavalli, Steven Seaman, Franz J Zemp, John Lee, Aly Abdelkareem, Katrina Ellestad, Alex Murison, Michelle M Kushida, Fiona J Coutinho, Yussanne Ma, Andrew J Mungall, Richard Moore, Marco A Marra, Michael D Taylor, Peter B Dirks, Trevor J Pugh, Sorana Morrissy, Bradley St Croix, Douglas J Mahoney, Mathieu Lupien, Marco Gallo

## Abstract

We investigated the role of 3D genome architecture in instructing functional properties of glioblastoma stem cells (GSCs) by generating the highest-resolution 3D genome maps to-date for this cancer. Integration of DNA contact maps with chromatin and transcriptional profiles identified specific mechanisms of gene regulation, including individual physical interactions between regulatory regions and their target genes. Residing in structurally conserved regions in GSCs was *CD276*, a gene known to play a role in immuno-modulation. We show that, unexpectedly, CD276 is part of a stemness network in GSCs and can be targeted with an antibody-drug conjugate to curb self-renewal, a key stemness property. Our results demonstrate that integrated structural genomics datasets can be employed to rationally identify therapeutic vulnerabilities in self-renewing cells.

**SIGNIFICANCE:** In adult GBM, GSCs act as therapy-resistant reservoirs to nucleate tumor recurrence. New therapeutic approaches that target these cell populations hold the potential of significantly improving patient care and overall prognosis for this always-lethal cancer. Our work describes new links between 3D genome architecture and stemness properties in GSCs. In particular, through integration of multiple genomics and structural genomics datasets, we found an unexpected connection between immune-related genes and self-renewal programs in GBM. Among these, we show that targeting CD276 with knockdown strategies or specific antibody-drug conjugates achieve suppression of self-renewal. Strategies to target CD276^+^ cells are currently in clinical trials for solid tumors. Our results indicate that CD276-targeting agents could be deployed in GBM to specifically target GSC populations.

**HIGHLIGHTS:**

- We generated high (sub-5 kb) resolution Hi-C maps for stem-like cells from GBM patients.
- Integration of Hi-C and genomics datasets dissects mechanisms of gene regulation.
- 3D genomes poise immune-related genes, including *CD276*, for expression.
- Targeting CD276 curbs self-renewal properties of GBM cells.

## INTRODUCTION

Glioblastoma (GBM) is the most common malignant brain tumor in adults and current treatments are mostly palliative (Stupp et al., 2005). Development of rationally-designed new treatments for this malignancy has been hampered by the high degree of intertumoral (Brennan et al., 2013; Verhaak et al., 2010) and intratumoral heterogeneity. Recent work has shown that intratumoral heterogeneity can be observed at the genomic, epigenomic and functional level. Genomic heterogeneity is manifested by the co-occurrence of multiple subclonal compartments in primary tumors and their dynamic selection in recurrences (Johnson et al., 2014; Meyer et al., 2015; Snuderl et al., 2011; Sottoriva et al., 2013; Szerlip et al., 2012). Functional heterogeneity is reflected by the non-equipotency of GBM cells: Only a subpopulation of cells with self-renewal properties contributes to sustained tumor growth. These self-renewing cells can engraft in transplantation experiments in immunocompromised mice and they contribute to genetic tumor evolution (Galli et al., 2004; Lan et al., 2017; Singh et al., 2004). We refer to these cells as cancer stem cells, or GBM stem cells (GSCs). GSCs are resistant to standard-of-care therapy and are thought to be responsible for tumor recurrence (Bao et al., 2006; Eramo et al., 2006). We and others have shown that chromatin (Gallo et al., 2012, 2015; Heddleston et al., 2012) and transcriptional profiles (Patel et al., 2014; Suvà et al., 2014) differ between GSCs and non-GSCs, suggesting that intratumoral epigenetic heterogeneity may reflect functional differences between populations of stem-like and more differentiated cells. It is therefore possible that specific epigenetic and chromatin states are pivotal in maintaining a functional hierarchy in GBM, similar to their roles in preserving hierarchical systems required for normal tissue homeostasis. In support of this hypothesis, several factors regulating the epigenome and chromatin states have been shown to be expressed predominantly in GSCs and to contribute to self-renewal properties, including EZH2 (Kim et al., 2013; Suvà et al., 2009), BMI1 (Abdouh et al., 2009; Jin et al., 2017), MLL1 (Gallo et al., 2012; Heddleston et al., 2012) and DOT1L (Miller et al., 2017). Some chromatin factors can also repress the self-renewing states. For instance, the histone variant H3.3 - a key nucleosome subunit - antagonizes self-renewal and is actively repressed in GSCs by an MLL5- mediated mechanism (Gallo et al., 2015). Electron spectroscopic imaging and ATAC-seq experiments have shown that repression of H3.3 in GSCs results in globally more "open" chromatin dotted by numerous foci of locally compacted chromatin. This chromatin conformation was drastically different from non-self-renewing GBM cells. These data point to important roles for chromatin architecture and epigenomic regulation in maintaining functional GSC properties. However, it is not currently known how chromatin states and cancer-associated genetic lesions interact to shape overall three-dimensional (3D) genome architecture in GSCs.

3D genome organization results from the stereotypical folding of each chromosome in three-dimensional space. A recently-developed technique called Hi-C (Lieberman-Aiden et al., 2009) allows investigations of genome architecture by measuring the frequency of physical interactions between genomic regions. Original work using Hi-C showed that the genome of embryonic stem cells is organized into regions about 1 Mb in size, termed topologically associated domains or contact domains (Dixon et al., 2012). DNA segments in each domain physically interact more frequently with each other than with DNA in other domains. Loop anchors form the boundaries of domains and are usually nucleated by the presence of anti-parallel binding sites for CCCTC-binding factor (CTCF) (Rao et al., 2014). CTCF and the cohesin complex are thought to bring distant genomic regions together by a loop-extrusion mechanism and play an essential role in insulating domains from each other (Ganji et al., 2018; Rao et al., 2017; Sanborn et al., 2015).

It was shown that domain boundaries act as barriers to the spread of chromatin compaction states, with each domain being predominantly in a compacted chromatin (repressed) state or in an open chromatin (active) state (Dixon et al., 2012). Interestingly, Hi-C contact maps show that domains with relatively open chromatin tend to interact with each other, while gene-poor and transcriptionally repressed regions tend to be sequestered at the nuclear lamina (Stevens et al., 2017). This broad-range separation of transcriptionally active and inactive regions defines type A (open chromatin) and type B (closed chromatin) compartments (Dixon et al., 2012; Lieberman-Aiden et al., 2009). While domains tend to be relatively conserved between cell types, the membership of each domain in type A or B compartments varies and may act as a barcode to identify a specific cell type (Dixon et al., 2015). CTCF is absolutely required for the formation and maintenance of domains, but it has no effect on the organization of compartments (Nora et al., 2017). Similarly, blocking the loading of cohesin on chromatin erases the configuration of domains, but it has minimal effects on genome compartmentalization (Schwarzer et al., 2017). These studies therefore suggest that CTCF and cohesin are required for setting up domain organization, whereas epigenetic and chromatin modifications instruct the overall organization of the genome into compartments.

Beside for domains and compartments, high-resolution Hi-C maps allow the visualization of individual loops that form between regulatory regions, such as enhancers and super enhancers (SEs) and their target gene. In addition to CTCF, loop formation can also be mediated by Yin Yang 1 (YY1) (Weintraub et al., 2017) and by ZNF143 (Bailey et al., 2015) in the context of enhancer-promoter interactions.

Thus, Hi-C data can provide different types of information: High-resolution data can identify precise DNA looping events; lower resolution data can identify self-interacting contact domains and longer-range chromosome compartmentalization (**Figure 1A**). Integration of information on loops, domains and compartment organization can provide a precise view of genome organization for specific cell types.

**Figure 1.**
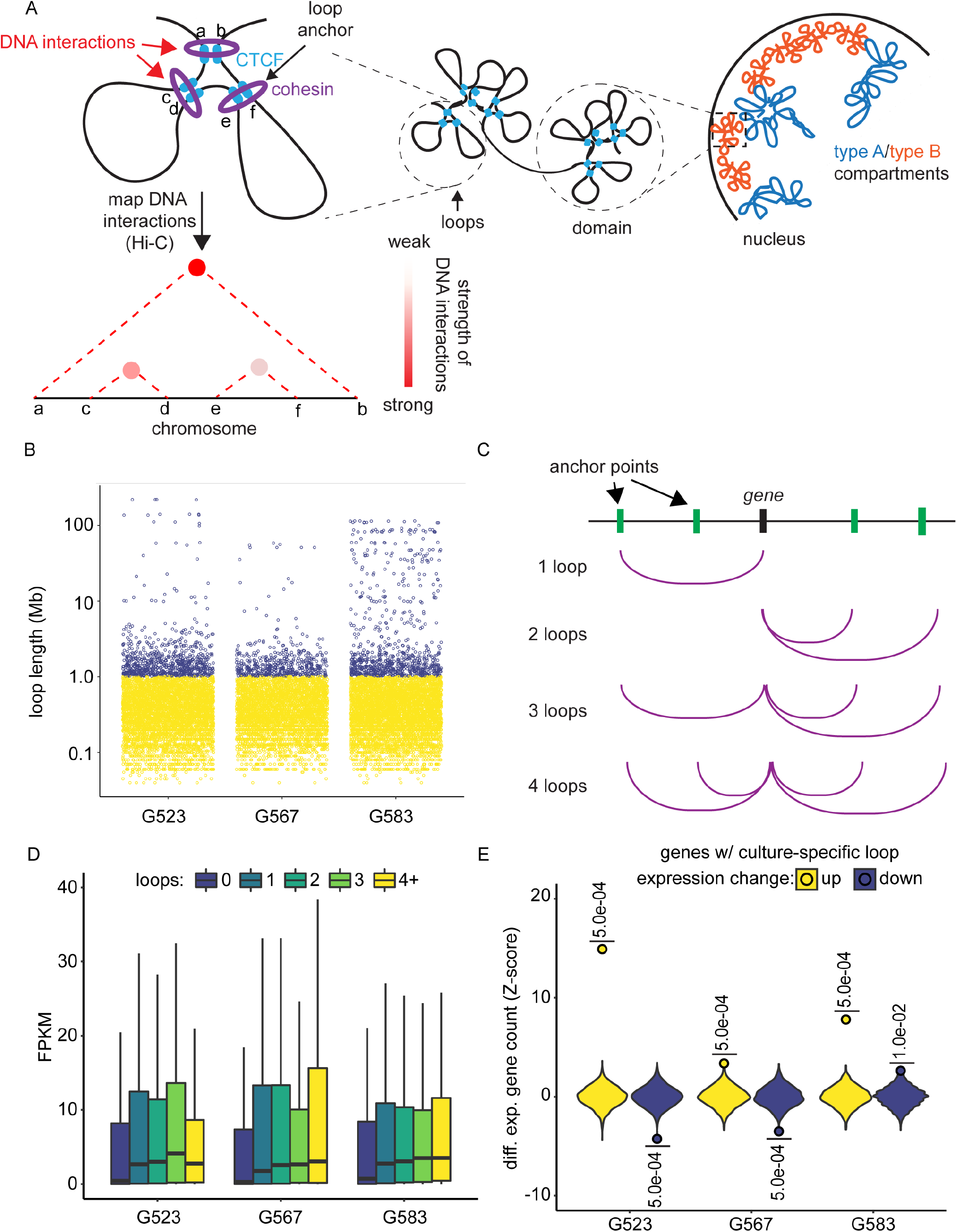
Loop architecture and its effect on gene expression in GSCs. (A) Diagram depicting hierarchical relationships 3D genome elements: Loops, domains and compartments. Hi-C measures the frequency of DNA interactions between DNA regions. (B) Distribution of loop lengths for three patient-derived GSC cultures (G523, G567 and G583). Yellow and blue colors were set at a 1 Mb threshold. (C) Cartoon representation of how a single gene may loop to multiple distinct control regions. The situations of a gene being engaged in 1, 2, 3 and 4 loops are shown here. (D) Genes engaged by at least one loop are more highly expressed than genes without loops. Expression trends upward with increasing numbers of loops. (E) Culture-specific loops are enriched for genes with elevated expression and depleted for genes with decreased expression. Dots: Measured count of differentially expressed genes found overlapping culture-specific loops. Violins: Distribution of 2000 counts of differentially expressed genes generated by randomly selecting expression values from genes with loops shared between cultures. See also Figure S1.

Recent work has shown that disruption of genome architecture may contribute to the etiology of developmental disorders (Lupiáñez et al., 2015) and cancer (Flavahan et al., 2016) through dysregulation of gene expression programs. Flavahan et al. provided evidence that hypermethylation of a CTCF binding motif in some *IDH1*-mutant gliomas eliminates a domain boundary and results in sustained expression of the *PDFGRA* oncogene. The global impact of genome architecture on transcriptional programs in gliomas and other brain tumors has been difficult to assess because of the lack of high-resolution 3D genome maps for these cancers.

We hypothesized that 3D genome architecture plays an important role in determining patient-specific transcriptional programs in GSCs. We have therefore generated high-resolution Hi-C maps of patient-derived primary GSC cultures. Integration of Hi-C, chromatin maps and transcriptional datasets for each sample generated a comprehensive view of the precise mechanisms of transcriptional control in GSCs at the level of individual genes. We provide proof-of-principle that 3D genome information can be used to identify new potential therapeutic targets to curb the self-renewal properties of GSCs.

## RESULTS

### High-resolution Hi-C maps identify long-range contacts in GSCs

To investigate 3D genome architecture in GBM, we generated DNA contact maps from low-passage (range: passage number 10-14) GSC cultures from three patients (G523, G567 and G583) by *in situ* Hi-C (Rao et al., 2014). We sequenced our Hi-C libraries to generate a total of 2.5 billion paired-end reads and achieved sub-5 kb resolution for each sample (see **Table S1** for quality control and characteristics of each library). To the best of our knowledge, these are the highest-resolution maps generated from patient-derived primary cultures. From this dataset, we identified loops, domains, and compartment organization for each sample (**Figures S1A-D**; **Tables S2-4**). We observed a wide range of loop lengths within each GSC culture. Over 80% of the loops were under 1 Mb in length (**Figures 1B and S1A,E**), which corresponds with the average length of contact domains in human cells (Dixon et al., 2012; Nora et al., 2012). However, we did observe many long-range contacts between regions separated by over 1 Mb, including rare loops over 100 Mb (**Figure 1B; Table S2**). Overall, loop calls between GSC cultures were largely conserved, with pairwise Spearman correlation scores ranging from 0.79-0.82 between cultures (**Figure S1B**). The concordance in compartment calls between GSC cultures was lower, with Spearman correlation scores in the 0.57-0.73 range (**Figure S1D**), suggesting a higher degree of dissimilarity at the compartment level than at the loop level.

DNA loop formation is an important component of transcriptional regulation. SEs can loop over large distances to contact the promoter regions of the genes they regulate. Although two-thirds of annotated genes (16,466 out of 24,261 predicted genes) had no detectable looping in any sample, we identified 3,111 genes engaged in at least two loops in any one GSC culture (**File S1**). We asked whether the number of loops that contact a single gene correlated with increased transcription. Integrating the loop calls from our Hi-C maps with RNA-seq data, we compared the expression levels of genes that did not engage in looping with genes that were engaged by one or more loops at the population level (**Figure 1C**). Genes that were contacted by a single loop were expressed at significantly higher levels than those that were not part of any loop (**Figure 1D**; p = 1.06 × 10^-61^ for G523, p = 6.18 × 10^-60^ for G567 and p = 4.03 × 10^-51^ for G583). Although the median expression value trended higher with increasing numbers of loops, there was no statistical difference between genes contacted by one loop and genes contacted by more than one loop (**Figure 1D**). These data suggest that at the population level, formation of a loop is necessary for gene expression, and additional loops converging on a gene have minimal effects on gene expression.

### Integration of Hi-C and chromatin datasets allow precise visualization of SE-gene interactions

SEs are large conglomerates of regulatory regions that can boost expression of their target genes to higher levels than regular enhancer elements (Whyte et al., 2013). Linking regulatory regions, including SEs, to the specific genes they target is a notoriously imprecise endeavour. Associating an SE to a gene immediately downstream simply based on proximity is often done, although this is an obviously imprecise approach. Alternatively, biochemical and reporter assays can be employed to validate the interaction between an SE and its putative target gene. We tested the ability of our Hi-C maps to detect SE-gene interactions. For each culture, we performed chromatin immunoprecipitation with massively parallel sequencing (ChIP-seq) for histone 3 lysine 27 acetylation (H3K27ac), which is a robust marker of open chromatin at enhancer, SE and promoter regions. Subsequently, we used these H3K27ac ChIP-seq datasets to call putative SEs with the ROSE package (Lovén et al., 2013; Whyte et al., 2013). As expected, genes targeted by SE-promoter loops had the highest level of expression, genes targeted by loops that did not involve SEs had lower expression levels, and genes with no loops had the lowest expression (**Figure S1F**). We also observed low correlation for H3K27ac peaks between samples, indicative of globally distinct transcriptional activity and enhancer usage in different GSC cultures (**Figure S1G**). These data show that our Hi-C maps have sufficient resolution to link SEs and other regulatory regions to individual genes.

### Low levels of conservation in CTCF occupancy between GSC samples

Although convergent CTCF motifs are nearly always found at loop anchors, there remain many predicted CTCF motifs in the genome that are not associated with loop formation. To precisely identify CTCF motifs involved in loop formation, we generated ChIP-seq datasets for CTCF for each GSC culture. Analysis of our ChIP-seq datasets showed relatively low conservation of CTCF binding across our GBM samples (**Figure S1H**). Overall, we identified the CTCF motifs utilized at both loop anchors for ~30% of loops, and a least one anchoring CTCF motif was identified for ~80% of loops (**Table S2**). These numbers are in strong agreement with high-resolution Hi-C data published for embryonic stem cells and other experimental systems (Dixon et al., 2012; Rao et al., 2014). A recently published report showed that hyper-methylation of CTCF binding sites can lead to suppression of CTCF binding (Flavahan et al., 2016). We therefore studied DNA methylation patterns in G523 and G583 with EPIC DNA methylation arrays. Of ~860,000 EPIC probes, we detected high methylation (β values > 0.8) at ~280,000 sites. Permutation analysis revealed that the number of methylated probes that overlap predicted CTCF sites was significantly lower than expected by randomly sampling the complete probe set (**Figure S1I**). Of the CTCF motifs utilized within loop anchors, only two motifs appeared methylated in either G523 or G583 (**Figures S1J**). Our data show that *IDH1*-wild type GBM may be less prone to undergo CTCF-dependent disruption of loop and domain organization than *IDH1*-mutant gliomas (Flavahan et al., 2016).

### Sample-specific loops drive differential regulation of gene expression

Although there was a strong correlation in looping between our samples (**Figure S1B**), there were also many loops that differed in their position and strength (**Figure S2A**). Position was defined as utilization of specific loop anchors ± 5 kbp to account for the resolution of our maps. Strength was defined as the number of paired-end reads in support of a specific interaction. We defined “private loops” as those present only in a single sample and asked whether private loops had measurable effects on transcriptional output. We identified private loops for each patient and tested for inter-sample differential expression of genes at the anchors of these loops using RNA-seq datasets. Permutation analysis indicated that the genes contacted by private loops were significantly enriched for elevated expression (**Figure 1E**). For G523 and G567, we also found depletion of genes with reduced expression. These results indicate that private loops play a significant role in enhancing expression of specific genes in each GSC culture.

One representative example is offered by the *QKI* locus, which differentially engages SEs in each of the three GSC cultures we profiled. The number of SEs looped to *QKI* in G523, G567, and G583 was three, zero, and two, respectively (**Figure 2A**). In addition, G523 had two private loops at the *QKI* locus (colored in light blue in **Figure 2A**). It is generally accepted that Hi-C can only identify up to two contacts for any given site per cell (one for each homologous chromosome in a diploid genome). However, a recent study showed that it is possible to detect multi-loop hubs occurring in individual cells (Rao et al., 2017). We investigated this possibility for *QKI*. We specifically looked for split sequencing reads that could map to more than two DNA regions. This analysis provides evidence for triplets of loop anchors, in different combinations, simultaneously interacting with each other (**Figure 2B**). This is direct evidence that *QKI* is in direct contact with more than one control region at the level of individual cells. We did not detect more than three loop anchors interacting at the same time, probably because of the limits imposed by read length during the sequencing process. In agreement with these data, G523 expressed almost 12 times more *QKI* transcripts as G567 and G583 (**Figure 2C**). These results show that although the total number of loops interacting with a gene is not proportional to expression levels (see above), private loops have a strong role in potentiating transcriptional output.

**Figure 2.**
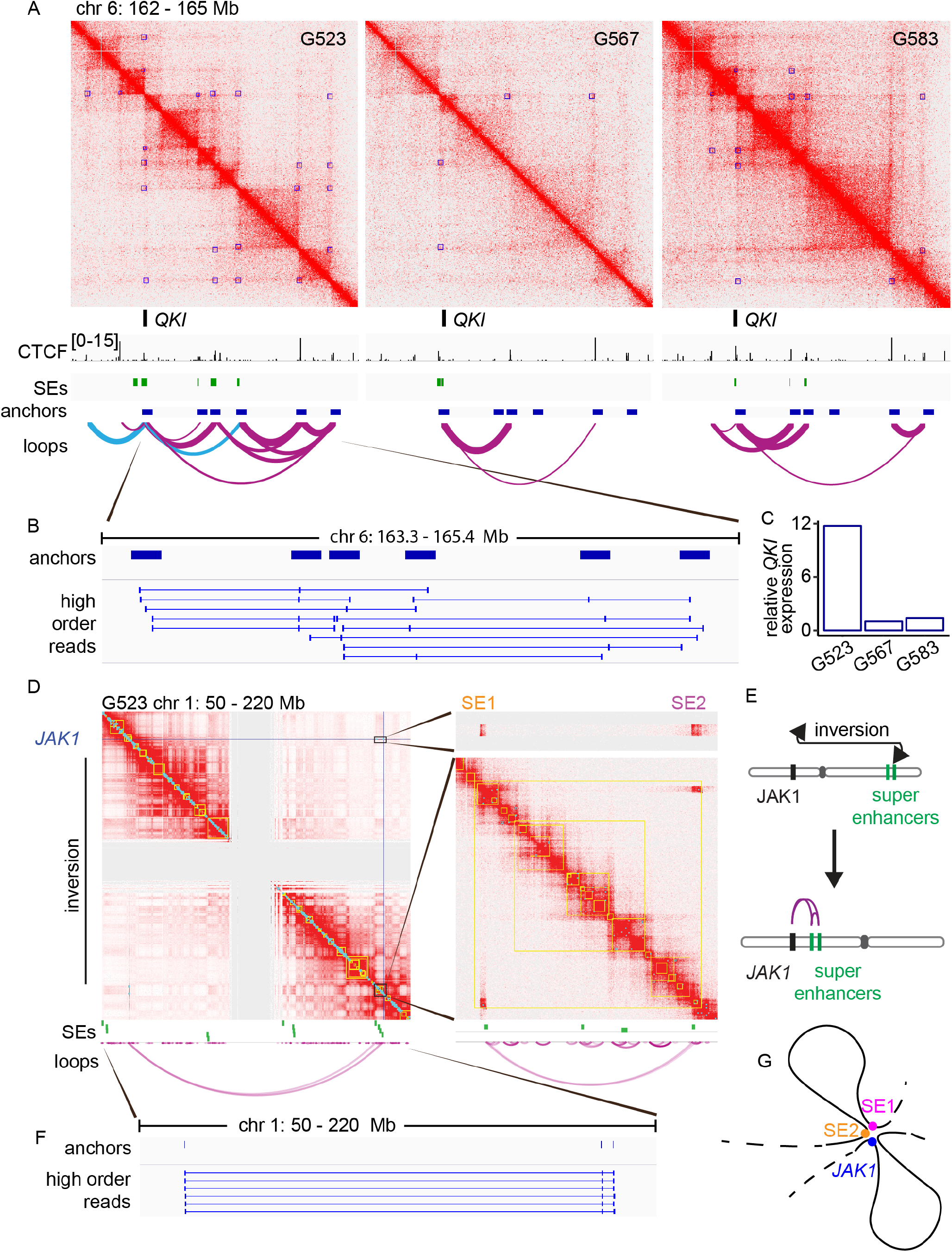
High-order loops and genomic rearrangements cause differential super enhancer interactions by GSCs. (A) Hi-C contact maps for G523, G567 and G583 surrounding the *QKI* locus. Super enhancers (SEs) were called using ROSE with H3K27ac ChIP-seq data. Loops identified with Hi-C data are shown in purple. Loops colored in light blue represent private loops for G523. (B) Chimeric reads derived from the same DNA fragment that also align to more than two loop anchors. (C) Expression of *QKI* in G523, G567 and G583 was determined by RNA-Seq. Y-axis represents read counts normalized to G567 to give fold enrichment values. (D) Hi-C contact maps assuming a standard chromosomal order indicate the formation of a ~140 Mb loop connecting *JAK1* to two super-enhancers at the other end of chromosome 1. The central grey region lacks signal throughout due to repetitive pericentromeric regions with ambiguous sequence alignments. (E) Schematic indicating how a large inversion brings the super enhancers and *JAK1* in close proximity. (F) Chimeric reads aligning to *JAK1* and both super-enhancers. Additional higher-order reads were detected but not all are displayed due to redundancy. (G) Diagrammatic representation of the convergence of two SEs (SE1 and SE2) to the *JAK1* locus in G523, as determined by Hi-C data. See also Figure S2.

### Structural variants can result in spurious engagement of SEs

We next asked whether large structural rearrangements may have an effect on 3D genome architecture, and more specifically on loops. Using whole-genome sequencing (WGS), we identified a large inversion on chromosome 1 that brought two SEs normally located on the q arm close to *JAK1* on the p arm in G523 (**Figures 2D,E**). Hi-C data showed clear physical interactions between the two SEs (SE1 and SE2) and *JAK1*, as illustrated by the heatmaps and the loop arc plots (**Figure 2D**). These interactions were not identified in the other two GSC cultures, which did not carry the chromosome 1 inversion. Our computational analysis found high order reads mapping to SE1, SE2 and *JAK1* (**Figure 2F**), providing evidence that these three elements form a contact hub in at least a subpopulation of cells (**Figure 2G**). This example illustrates that new SE-gene interactions can emerge as a consequence of structural variation in cancer cells. This is a particularly interesting example, because *JAK1* is known to play roles in GBM (McFarland et al., 2011) and other cancer types (Thomas et al., 2015).

### Genes connected to multiple loops are enriched for metabolic, stemness and cancer genes

Next, we investigated the distribution of loops per gene in our datasets. By looking specifically at genes that were targeted by at least two loops in any one of the samples analyzed by Hi-C, we identified a list of 3,111 genes (**File S1**). We ranked these genes based on the maximum number of loops they were associated with in any one GSC culture. Of these genes with > 2 loops, 97% were targeted by 5 or fewer loops. 3% of these genes (94 of 3,111 genes) displayed extreme interactivity, engaging with 6 or more loops. *VSNL1* was targeted by 23 loops, the highest number in our dataset (**Figure 3A**). This gene encodes a neuronal calcium sensor and is highly expressed in cerebellar granule cells (Bernstein et al., 1999; Braunewell et al., 2001). The second-highest number of loops interacted with *ALKBH5*, a gene encoding an m^6^A demethylase recently shown to have a role in GBM (Zhang et al., 2017). Zhang et al showed that ALKBH5 contributes to the tumorigenic properties of GSCs by stabilizing the *FOXM1* transcripts, resulting in increased expression for this gene. Furthermore, high *ALKBH5* expression is a poor prognostic factor for GBM patients. *IGFBP2* was targeted by 18 loops, and this gene plays an important role in regulating insulin-like growth factor signaling and the stemness properties of GSCs (Hsieh et al., 2010). *ANK2* was targeted by 10 loops. This gene was previously shown to be positively regulated by SOX2 (Berezovsky et al., 2014), which imparts important self-renewal and tumor-initiating properties to GSCs (Gangemi et al., 2009). *SOX2-OT*, a large gene that overlaps with *SOX2*, was also among the genes with the most connections.

**Figure 3.**
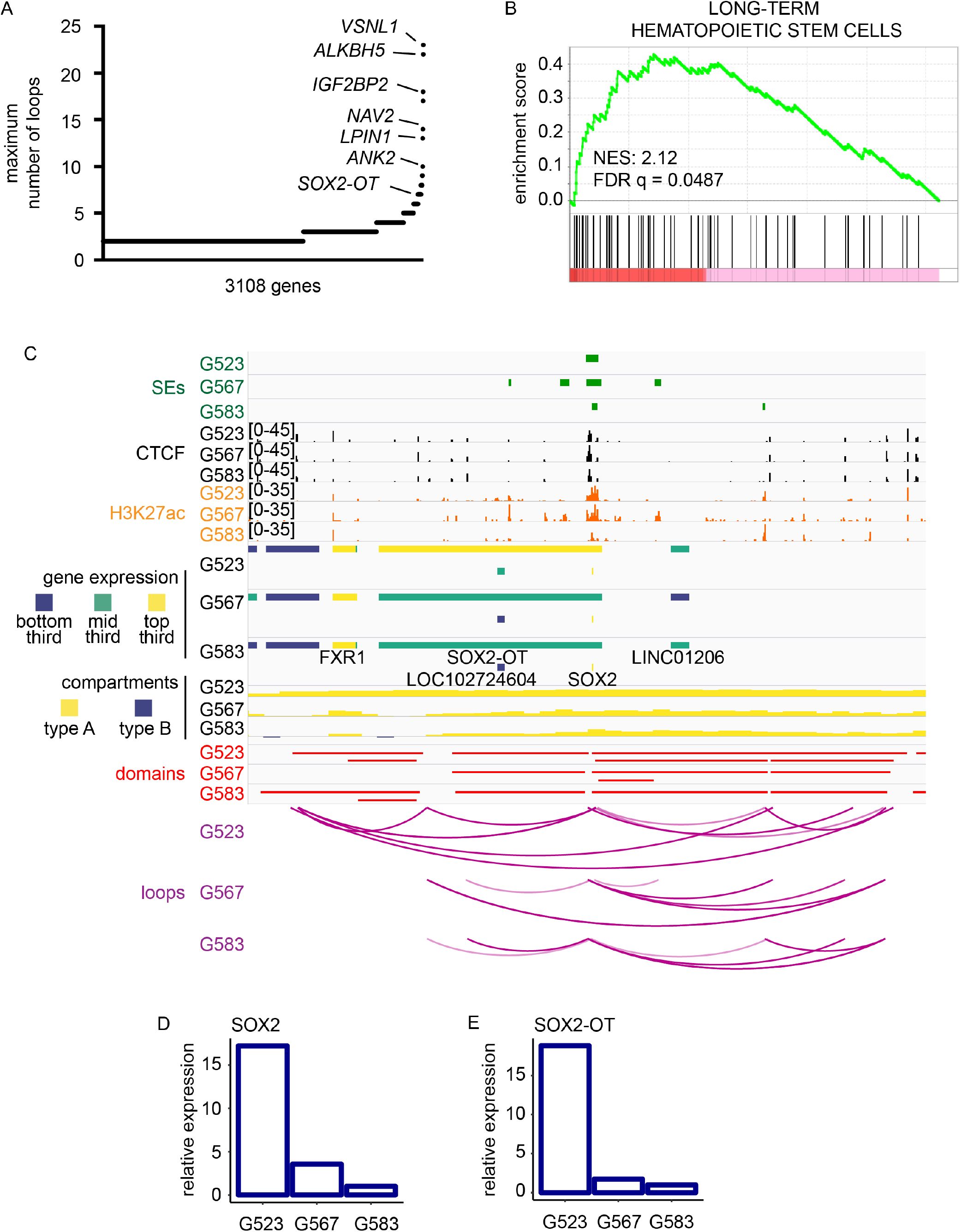
Interplay of 3D genome organization and chromatin features in transcriptional control of stemness genes in GBM. (A) 3108 genes were ordered based to the number of loops they contact. Each gene represented is part of at least two loops. (B) Gene set enrichment analysis of genes in (A) ranked based on the number of loops they contact. (C) Integration of genome browser tracks for ROSE super-enhancer calls, CTCF ChIP-seq, H3K27ac ChIP-seq, RNA-seq, compartments, domains, and loops, and chimeric reads determined by Hi-C at the *SOX2* locus. (D, E) Relative expression of *SOX2* and *SOX2-OT* as determined by RNA-Seq. See also Figure S3.

Given that some of these highly connected genes had established roles in GSC maintenance, we ranked this set of genes by the number of loops engaged and performed gene set enrichment analysis (GSEA) (Subramanian et al., 2005). We found significant positive enrichment of genes involved in regulation of insulin secretion (normalized enrichment score (NES): 2.19; q = 0.0886) (**Figure S2B**) and metastasis (NES: 2.16; q = 0.0530) (**Figure S2C**). Interestingly, we also found enrichment for genes associated with long-term hematopoietic stem cells (Ivanova et al., 2002) (NES: 2.12; q = 0.0487) (**Figure 3B**). Overall, our data show that highly connected gene sets in GSCs are significantly enriched for genes with established roles in cancer and in regulation of stemness properties.

### Integration of Hi-C and chromatin data allows dissection of mechanisms of transcriptional regulation

Expression of stemness genes is important for maintenance of GSC self-renewal, a key cancer stem cell property (Al-Hajj and Clarke, 2004; Laks et al., 2009). Some of these stemness factors are expressed at surprisingly different levels in GSCs from different patients. One example is provided by the stemness gene *SOX2*. At the *SOX2* locus, we find similar compartment and domain organization between GSC samples. However, there are several G523 private loops emanating from an upstream region (**Figure 3C**). Levels of H3K27ac at *SOX2* are very similar in G523 and G567. However, at the transcriptional level, *SOX2* expression is much higher in G523 than G567 and G583 (**Figure 3D**). Transcription of the overlapping non-coding RNA *SOX2-OT* was also elevated in G523 compared to the other samples (**Figure 3E**), as this gene and *SOX2* might share regulatory regions and mechanisms. High order reads indicate that multiple loops simultaneously contact *SOX2* in individual cells to boost its expression in G523 (**Figure S3A**).

Another example is provided by *ASCL1*, which was shown to be important for the maintenance of self-renewal and tumorigenic properties of GSCs (Rheinbay et al., 2013; Suvà et al., 2014). However, *ASCL1* is expressed at high levels in some patient-derived GSCs, and at very low levels in others (Park et al., 2017; Suvà et al., 2014). The reason for these differences in *ASCL1* expression in different subgroups of GSCs is not currently known. Our Hi-C data, ChIP-seq and transcriptional profiles of GSC cultures gave us an opportunity to infer mechanisms of regulation of this important stemness gene. Our Hi-C data show that the overall domain organization and looping patterns at the *ASCL1* locus is conserved between samples (**Figure S3B**). However, expression of this gene is ~50-fold higher in G523 than in G567, and ~5-fold higher in G523 than in G583 (**Figures S3C**). Interestingly, *ASCL1* is in a type A compartment in G523 and type B compartment in G567 and G583. H3K27ac signals are observed only in G523 (**Figure S3B**). These findings suggest that loops and domains alone do not mandate transcription, but chromatin states play a strong role. These data are in agreement with recent evidence that 3D genome architecture changes temporally precede chromatin mark deposition that affect transcription during reprogramming of somatic cells into pluripotent stem cells (Stadhouders et al., 2018).

These examples highlight the multi-layered regulation associated with differential genome architecture. While it is likely that loops, domains, and compartments are each independently insufficient to predict the activation of a particular gene, combining these data types with maps of chromatin marks can explain differential transcriptional patterns observed between otherwise similar GSC cultures. Integration of 3D genome architecture and chromatin marks may therefore be necessary for fine tuning transcriptional activity.

### Discordant genome compartmentalization between GSC cultures

Domain lengths were uniform across samples, even after calling domains at different resolutions (**Figure 4A**). Length distributions and total numbers of compartment calls were also uniform across GSC cultures (**Table S3**).

**Figure 4.**
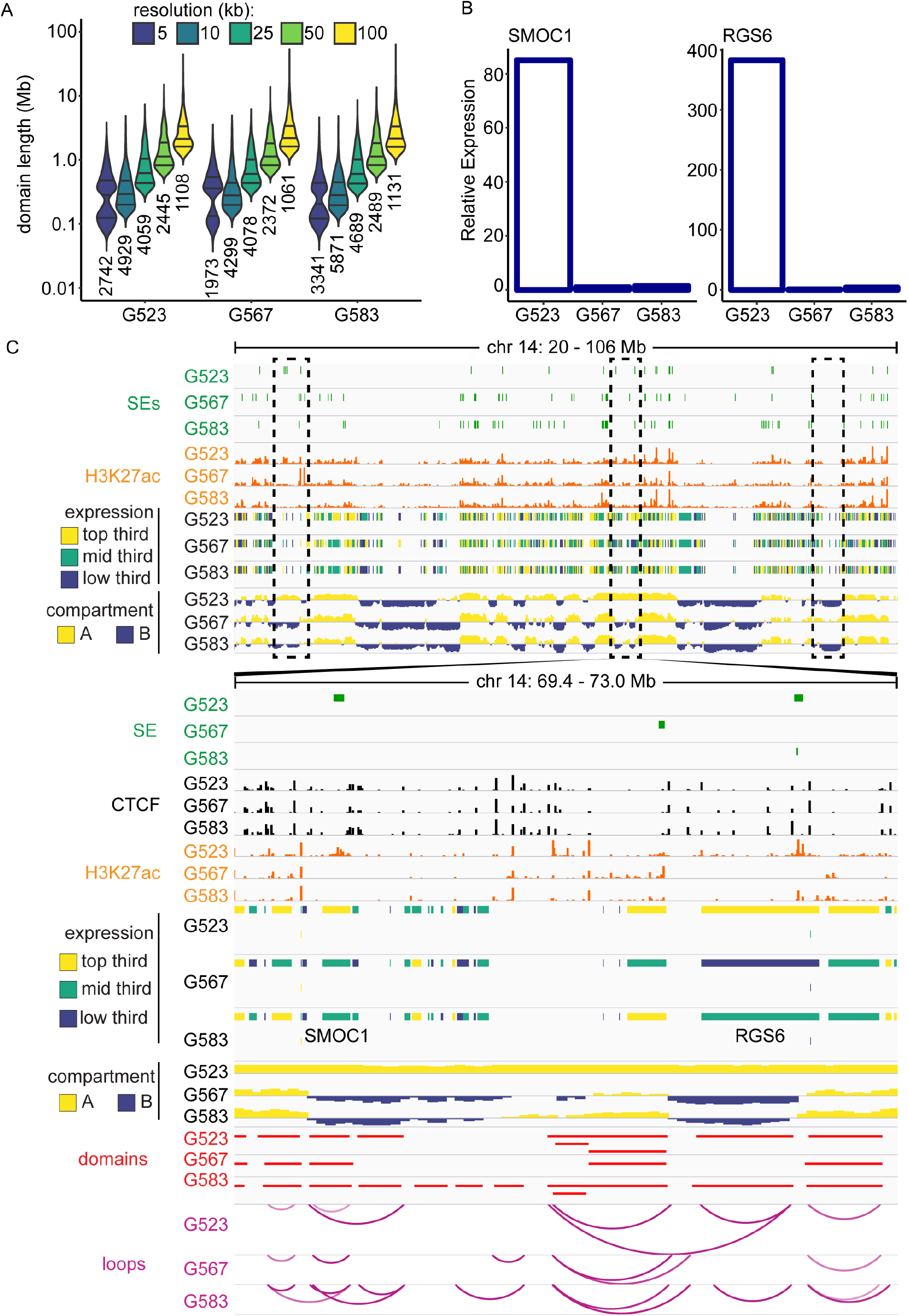
Structure of domains and compartments in GSC cultures. (A) Lengths of topological domain in GSC cultures. Domains were called at different resolutions. Total number of domains called at each resolution is on top of each violin plot. Horizontal lines in each violin plot represent quartiles. (B) Expression of *SMOC1* and *RGS6*, identified as differentially expressed genes with culture-specific compartmentalization. (C) Example of culture-specific compartmentalization in GSCs. For CTCF ChIP-seq data, tracks are scaled at [0-20] reads per million total reads. Tracks for H3K27ac ChIP-seq are scaled to [0-25] reads per million total reads. See also Figure S4.

Using our Hi-C data, we called type A compartments (characterized by open chromatin and higher transcriptional levels) and type B compartments (characterized by compacted chromatin and lower transcriptional levels) for all three GSC cultures. We then used our H3K27ac ChIP-seq data to corroborate compartment assignment. We were able to define compartment status for 93% of the genome (2.86 Gbp of 3.09 Gbp in genome version hg38) in at least one culture. Similar numbers and sizes of compartments were identified in each culture. (**Table S4; File S3**). We found that genes in type A compartments exhibited elevated expression while genes in type B compartments were repressed (**Figure S4A**), as expected, providing further validation of our Hi-C data. In fact, the median expression of genes in type B compartments was zero (**Figure S4B**), indicating significant repression of transcription in these regions.

Next, we sought to identify regions of type A/B compartment similarity or dissimilarity between GSC cultures. Overall, compartment calls between the cultures were weakly but positively correlated (**Figure S1D**). Only 53% (1.52 Gbp of 2.86 Gbp definable) were called as type A or type B in all three cultures. The remaining 47% of the genome differed in its compartmentalization state between the three cultures. These data indicate that although there are similarities in compartmentalization between GSC samples, whether a given compartment is type A or B varies considerably between GSC cultures.

Accordingly, we identified regions of culture-specific compartmentalization – regions where compartmentalization was assigned in all three cultures, but compartment type was discordant between one GSC culture and the other two. Genes within private compartment regions were significantly more likely to be differentially expressed between cultures than by chance alone (**Figure S4C**).

We present the region of chromosome 14 containing *SMOC1* and *RGS6* as a representative example of differentially regulated genes between GSC cultures. *SMOC1* was expressed ~80-fold higher in G523 than in the other GSC cultures, and *RGS6* was expressed ~400-fold higher in G523 than in the other two cultures (**Figure 4B**). In this region we observe type A compartmentalization only in G523, as well as subtle differences in loops and domain structures between cultures (**Figure 4C**). Of note, H3K27ac signal was present at the *RGS6* locus in both G523 and G583, although the transcriptional output was radically different in these cultures. Looping architecture also differs at this locus between the two cultures. These data highlight the regulatory impact of genome compartmentalization and loops.

### Consensus type A compartments in GSCs are enriched for immune-related genes

We next asked whether we could identify regions of the genome with conserved compartmentalization among the three GBM cultures. As a non-GBM control we used previously published Hi-C data generated with GM12878, an immortalized lymphoblastoid cell line, because Hi-C contact maps of sufficiently high resolution to be comparable to ours are available for this line (Rao et al., 2014). We identified 3,021 genes whose compartmentalization was conserved among GSC samples but differed from GM12878 (**File S2**). Focusing on genes in GSC-specific type A compartments, we ranked the genes by expression, then used this ranked list for GSEA against datasets in the MSigDB collection. The data show enrichment for immune response (module 75) (**Figure 5A**) and thymus genes (module 44) (**Figure S5A**). Both gene sets are enriched for immune-related genes. This was an unexpected finding, but it suggested that GSCs might have mechanisms to express specific immuno-modulatory genes to escape immune detection. We decided to follow up on this finding by looking for candidates that could potentially be targeted in GSCs. Specifically, we looked for immune-related genes that met the following criteria: (a) located in GBM-specific type A compartments; (b) highly expressed in GSC cultures; (c) encoding cell surface proteins; (d) potentially targetable with current or experimental compounds. The immune-related gene *CD276* (also known as B7-H3) met all these criteria.

**Figure 5.**
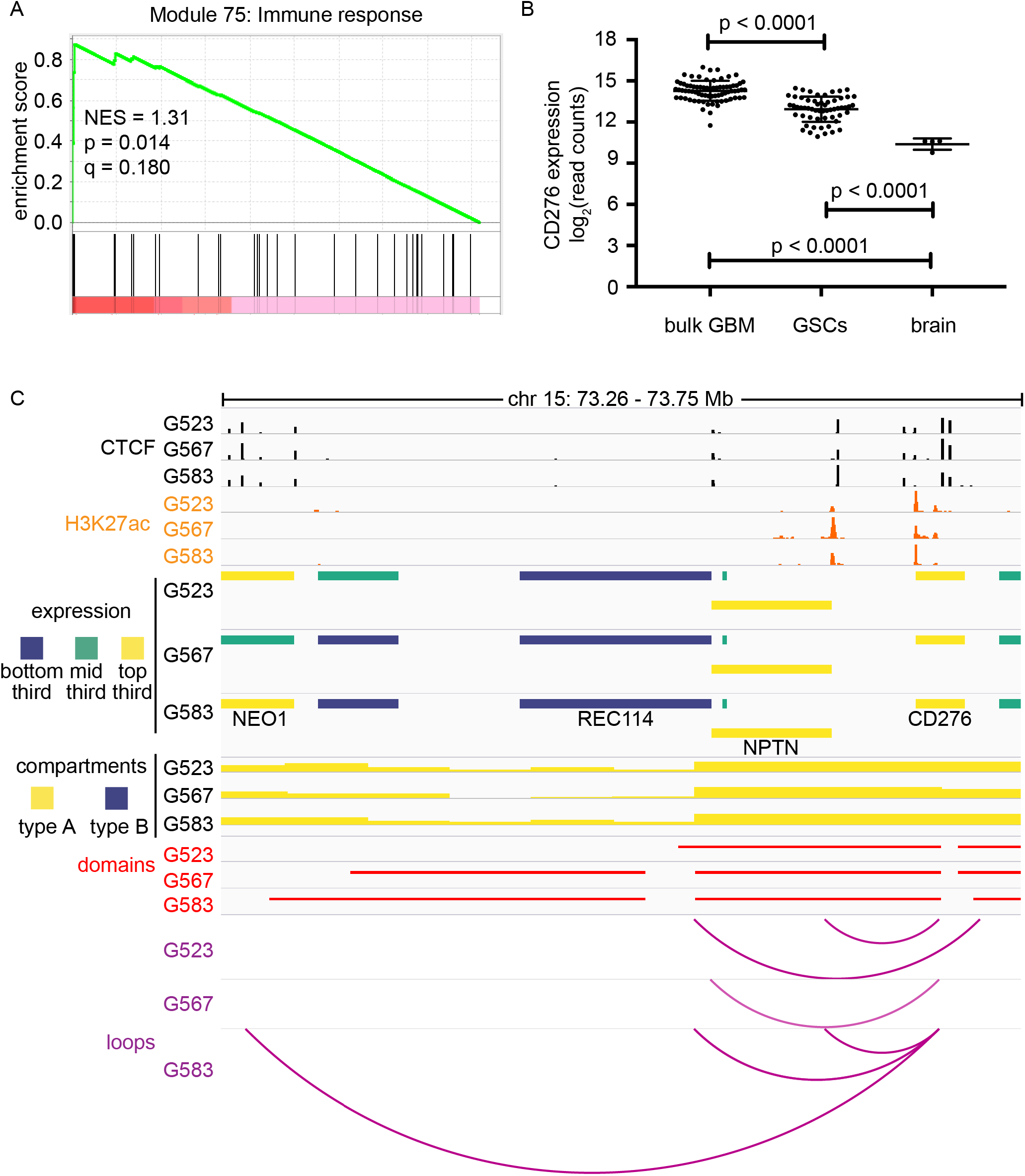
CD276 is an immune gene associated with stemness in GSCs. (A) GSEA was performed on the set of genes located in type A compartments in all GBM samples, but not in control GM12878 cells. The analysis identified significant enrichment in immune response genes. NES: Normalized enrichment score. (B) Expression of *CD276* was determined by RNA-seq in bulk GBM samples (n = 76), GSCs (n = 76) and non-neoplastic brain tissue (n = 4). P-values were calculated with the Mann-Whitney statistical test. (C) 3D genome and chromatin landscape at the *CD276* locus. This panel integrates H3K27ac ChIP-seq data, CTCF ChIP-seq data, 3D genome features called using Hi-C data (domains, compartments and loops), and RNA-seq data (gene expression) for three patient-derived GSC cultures (G523, G567 and G583). See also Figure S5.

### *CD276* is highly expressed in patient-derived GSCs

CD276 is a putative immune checkpoint (Lee et al., 2017) and is thought to be a repressor of T cell activation, preventing release of IFN-γ by T cells and impairing CD4^+^ cell proliferation (reviewed in Picarda et al., 2016). However, its exact role in immunomodulation is still controversial and appears highly dependent on the experimental models used (Hofmeyer et al., 2008). *CD276* is aberrantly expressed in a number of solid tumours and is detected on both tumour cells and in the tumour vasculature (Seaman et al., 2017). We found that *CD276* expression was higher in GSC cultures (n = 76) than in bulk GBM samples (n = 76; p < 0.001) or non-neoplastic brain tissue (n = 4; p = 0.0005) (**Figure 5B**).

In all three GSC cultures profiled by Hi-C, the *CD276* locus forms a CTCF-delimited loop ~140 kb upstream to the 5’ ends of genes *NPTN* and *REC114* (**Figure 5C**). In G523 and G583, loops connect a putative enhancer element upstream of *CD276* to the gene. In G583, we also detected a longer loop to CTCF binding sites within the gene body of *NEO1*. Therefore, although all three GSC cultures have high expression of *CD276* (top third of expression genome-wide, **Figure 5C**) and similar levels and profiles for H3K27ac, its transcriptional regulation might be finetuned by genome architecture in different samples. This is reflected by different transcriptional outputs for *CD276* in the three GSC cultures (**Figure S5B**).

Considering that the patient-derived primary cultures we used are enriched for self-renewing cells, we asked whether *CD276* expression correlates with expression of self-renewal genes. We also tested the hypothesis that *CD276* expression may correlate with expression of other immune genes, based on the GSEA data presented above. We calculated Pearson correlation scores for a previously published 16-gene GBM self-renewal signature (Suvà et al., 2014) and for a selected subset of immune genes that we found to be highly expressed in GSCs, using bulk RNA-seq data for our collection of patient-derived GSC cultures (n = 76). We found that although *CD276* expression most tightly clustered with other immune modulators (*IFNGR1, IFNGR2, TNFRSF1A* and *TNFSRF1B*), it was also positively correlated with expression of a cluster of self-renewal genes including *VAX2, SOX21*, and *CITED1* (**Figure 6A**). This association with the stemness signature was unique to *CD276*, as the other immune genes found were anticorrelated with the stemness signatures.

**Figure 6.**
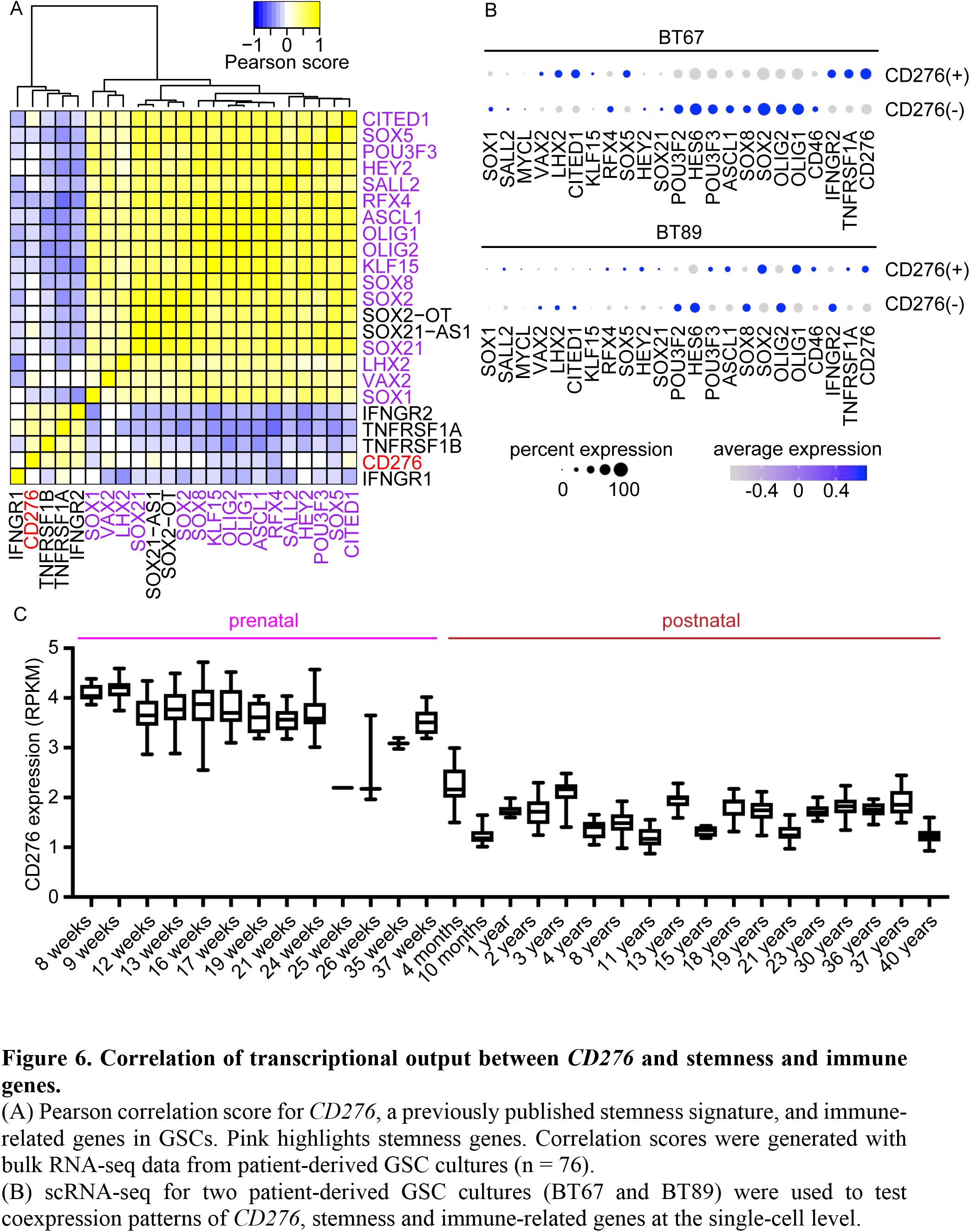
Correlation of transcriptional output between *CD276* and stemness and immune genes. (A) Pearson correlation score for *CD276*, a previously published stemness signature, and immune-related genes in GSCs. Pink highlights stemness genes. Correlation scores were generated with bulk RNA-seq data from patient-derived GSC cultures (n = 76). (B) scRNA-seq for two patient-derived GSC cultures (BT67 and BT89) were used to test coexpression patterns of *CD276*, stemness and immune-related genes at the single-cell level. (C) Patterns of expression of *CD276* in the prenatal and postnatal human brain. Data were extracted from Brainspan. See also Figure S6.

To deconvolute the correlation between *CD276* and the expression of self-renewal and immune-related genes in bulk RNA-seq datasets, we performed single-cell RNA-seq (scRNA-seq) on GSCs from two patients. Both cultures showed heterogenous *CD276* expression (**Figures S6A,B**). Subgroup analysis of *CD276^+^* and *CD276^-^* cells showed that *CD276^+^* GSCs tended to co-express subsets of stemness genes (**Figure 6B**). While *CD276^+^* cells co-express the stemness gene *SOX5* in both cultures, we observed variations in the repertoire of co-expression for other stemness gene. For instance, *CD276*, *OLIG1* and *SOX2* tended to be co-expressed in the same cells in BT89, whereas *CD276* tended to be co-expressed with *CITED1*, *LHX2* and *VAX2* in BT67. These data therefore suggest that *CD276* might be associated with a self-renewal program, although this program may be dependent on different subsets on stemness genes in the different samples.

Intriguingly, our meta-analysis shows that *CD276* expression in the human brain is higher during the prenatal period (average RPKM 3.74) than the postnatal years (average RPKM 1.72, two-tailed t-test p = 1.04 × 10^-211^; **Figure 6C**). This immune-related gene might therefore be associated with a primordial developmental program, providing a rationale for its association with the expression of self-renewal genes in GSCs.

### Targeting CD276 reduces self-renewal properties of GSCs

Given the link between *CD276* and the expression of stemness genes in GSCs, we explored further a possible role for this gene in the self-renewal program of GBM cells. We performed an *in vitro* assay to curb self-renewal in GSCs using an established growth factor withdrawal protocol (Gallo et al., 2015; Pollard et al., 2009). Growth factor withdrawal resulted in decreased protein levels for CD276 in G523 and G583 (**Figure 7A**), although it had no effect on G567 (**Figure S7A**). In a complementary experiment, we asked whether reducing levels of CD276 resulted in decreased self-renewal. To answer this question, we knocked down *CD276* using two inducible shRNA constructs (sh-CD276a and shCD276b) (**Figure 7B**). The effects of *CD276* knockdown on self-renewal were assessed with *in vitro* limiting dilution assays (LDAs). For both G523 and G583, knockdown of *CD276* resulted in reduction of self-renewal (**Figure 7C**).

**Figure 7.**
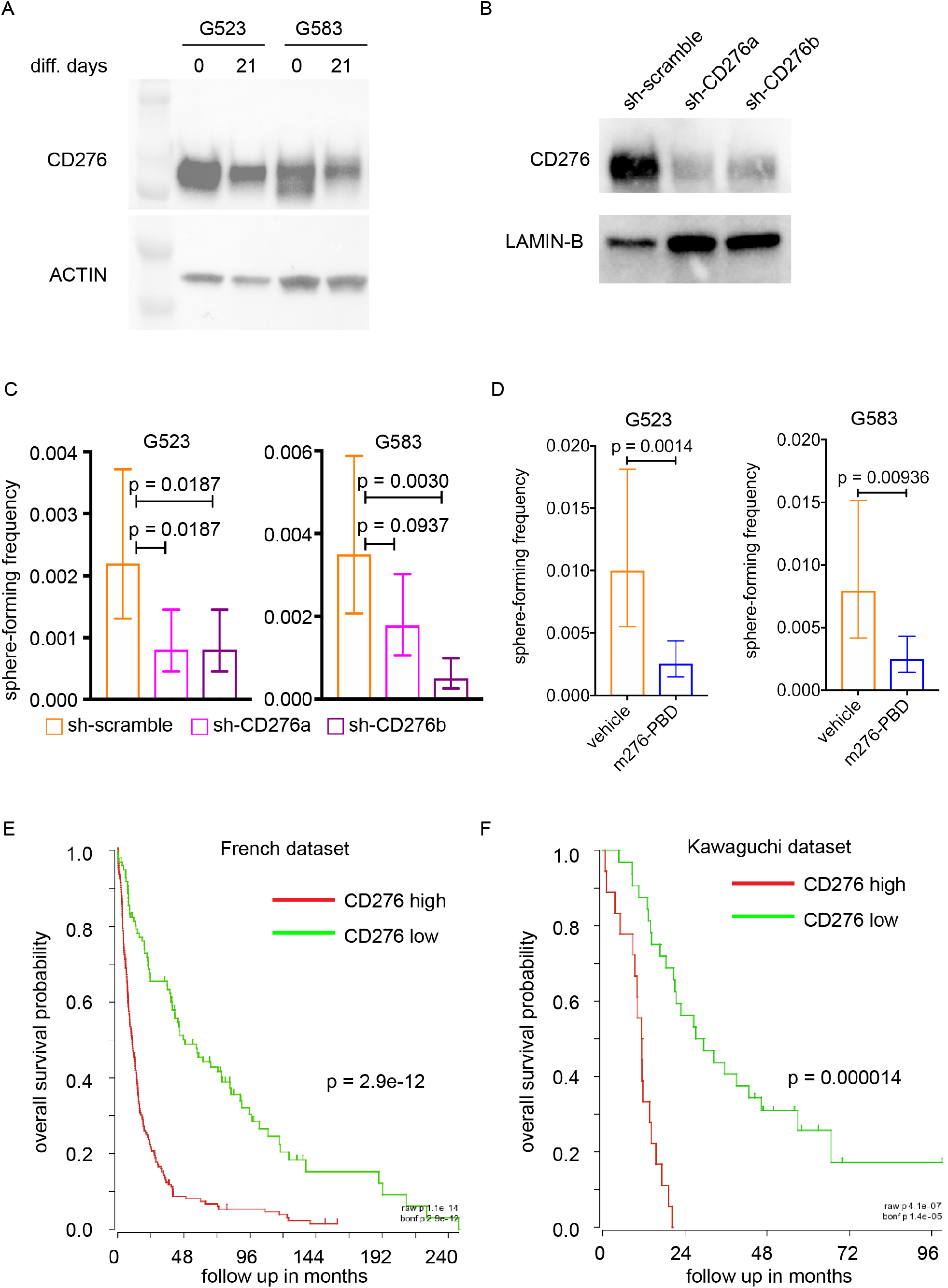
Targeting CD276 with an antibody-drug conjugate decreases self-renewal of GBM cells. (A-B) Western blots comparing CD276 levels between cultures exposed to (A) growth factor withdrawal or (B) shRNA constructs targeting *CD276*. (C) Limiting dilution analysis for G523 and G583 transfected with either scramble (control) or shRNA constructs targeting *CD276* (sh-CD276a and sh-CD276b). Data show mean sphere-forming frequency. Error bars correspond to 95% confidence interval. P-values were determined with ELDA. (D) Limiting dilution analysis for G523 and G583 treated with m276-PBD or vehicle control. Data show mean sphere-forming frequency. Error bars correspond to 95% confidence interval. P-values were determined with ELDA. (E-F) Survival of glioma patients stratified by *CD276* expression in the (E) French or (F) Kawaguchi datasets. Median gene expression was used to stratify patients. P-value was derived with log-rank statistics. See also Figure S7.

Considering that reduction of CD276 levels caused reduced self-renewal properties in GSCs, we explored tools that allow targeting CD276^+^ cells and have translational potential. Therapeutics against CD276, including a bispecific CD3-CD276 antibody (ClinicalTrials.gov identifier NCT02628535), are active agents used in clinical trials. We explored an alternative strategy, using an anti-CD276 monoclonal antibody conjugated with pyrrolobenzodiazepine (PBD) to selectively eradicate CD276 positive cells. This m276-PBD antibody-drug conjugate (ADC) was recently shown to be selective in delivering the PBD warhead to CD276^+^ cells in tumours and vasculature and to reduce tumor burden in several xenograft and mouse models of solid tumors (Seaman et al., 2017). We performed dose-response assays to determine concentrations of m276-PBD that were sub-lethal in these GSC cultures (**Figure S7B**). We then exposed G523 and G583 to IC_10_ concentrations of m276-PBD (see Methods) to assess the effects of the ADC on self-renewal in LDAs. For both G523 and G583, m276-PBD effectively curbed self-renewal (**Figure 7D**). These data support a role for CD276 in a self-renewal program in GSCs and validate the use of strategies to target CD276^+^ cells in adult GBM. In addition, our meta-analysis of previously published datasets shows that expression of *CD276* stratifies adult glioma patients, with high *CD276* expression being a negative prognostic factor (Bonferroni-corrected log-rank p = 2.9 × 10^-12^ for French dataset, **Figure 7E**; p = 0.000014 for the Kawaguchi dataset, **Figure 7F**; log-rank p = 0.01494 for TCGA GBM dataset, **Figure S7C**; p = 0.002094 for TCGA low grade glioma dataset, **Figure S7D**). All together, these data strongly suggest that targeting CD276^+^ cells might be a promising strategy to curb cancer stem cell properties in adult gliomas.

## DISCUSSION

We report that inter-sample 3D genome heterogeneity is a molecular feature of GBM. This conclusion was made possible through the generation of Hi-C maps for patient-derived GSCs with the highest resolution (< 5 kbp) for this cancer. This level of resolution allowed the visualization of individual loops connecting regulatory regions and their target genes. Even when our data are considered in the context of 23 Hi-C datasets generated from established cell lines, only GM12878 has a higher resolution (1 kbp) (Rao et al., 2014; Sauerwald and Kingsford, 2018). Together with the other genomic datasets we generated, these Hi-C maps will be an important resource for the brain tumor field, and for future studies on the relationships between 3D genome, epigenetic factors and transcriptional regulation. All our datasets will be immediately available, with Hi-C data readily displayed in a usable format for most users on the WashU Epigenome Browser.

Because of the long-range activity of regulatory regions, associating the activities of a specific SE (or a set of SEs) to a gene is notoriously difficult. Most accepted paradigms link an SE to its immediately downstream gene. This approach might have validity in some cases, but every SE-gene call should be experimentally validated. Our high-resolution Hi-C maps provide information on individual physical contacts between regulatory regions and their cognate genes across several GSC cultures. Interestingly, we found some loops enhanced gene expression even though they did not overlap with either CTCF binding site, enhancers or SEs. These loop anchors might me binding sites for other loop-inducing proteins, like YY1 or ZNF143, for instance. The characterization of protein occupancy and chromatin marks at these sites in GSCs will be the focus of future investigation. The availability of high-resolution Hi-C maps for a variety of cell and tumor types will be fundamental to provide mechanistic insight into the transcriptional and epigenetic control of gene expression in healthy and disease states.

Our Hi-C maps highlighted two major principles relating to genome organization: Individual local loops and high-order compartmentalization are different between GSC cultures, whereas domain organization is largely conserved. These findings are largely consistent with the notion that about 80% of domains are conserved among cell lines derived from different tissues types (Dixon et al., 2012). This is probably because domain boundaries often coincide with DNA replication start sites (Pope et al., 2014) and their conservation may therefore be evolutionarily constrained. On the contrary, individual looping interactions and compartments seem shaped by transcription factor activity and chromatin landscapes, and therefore exhibit more variation. Importantly, although the total number of looping interactions with a specific gene was not correlated with increased expression, private loops had a strong effect on potentiating transcriptional output. Some of these private loops might stem for engagement of SEs that are not available in other samples, and might reflect mechanisms of epigenetic evolution of the cancer cells.

The differences we observed at the level of loops and compartments between GSC cultures indicate that transcription factor and epigenetic dynamics are the driving forces in specifying transcriptional programs. Consequently, loop and compartment differences can be used to infer how changes in transcriptional output are achieved between patient-derived samples. In this regard, loops and compartments that are conserved among GSC cultures may underlie epigenetic states and transcriptional programs that are fundamental for the maintenance of cancer stem cell properties, including self-renewal. Using primary surgical resections, we previously reported that the key self-renewal factors like SOX2 are widely expressed in GBM cells, with between 5% and 50% of cells being SOX2^+^ in different samples (Gallo et al., 2012). This result is in apparent contradiction with functional studies that showed that cancer stem cells are rare in this tumor type (Singh et al., 2004). However, our data show stemness genes tend to be coexpressed in GSCs, leading to choreographed transcriptional signatures of self-renewal. This choreography is achieved through the integration of epigenetic inputs and transcription factor occupancy and activity to achieve consensus 3D genome states in genomic regions containing key stemness genes. Stemness is achieved through co-regulation of many genes that participate in the self-renewal signature, and cannot be achieved via the expression of an individual gene. This idea is supported by the requirement for the overexpression of multiple key stemness transcription factors to successfully reprogram non-GSCs to the GSC state (Suvà et al., 2014). Based on data published over the last decade, GSCs are thought to give rise to progenitor-like cells with decreased self-renewal properties, which eventually produce non-self-renewing cells that contribute to tumor bulk and ultimately are destined to die. We therefore hypothesized that GSCs need to achieve specific chromatin states and 3D genome architecture to maintain their self-renewal transcriptional network. We expect that perturbation of these states will lead to destabilization of transcriptional programs of self-renewal with consequent loss of stemness properties. Identification of ways to induce loss of self-renewal properties is a key goal of research using GSCs, and could have important implications for patient outcomes. Given that self-renewing cells have been shown to be therapy-resistant (Bao et al., 2006), they are thought to nucleate recurrences after administration of standard of care. Killing GSCs or causing them to lose self-renewal properties could be an effective way of delaying or preventing tumor recurrence, which is the most common cause of death for GBM patients.

By looking for consensus 3D genome features among GSC cultures, we aimed at identifying new members of the self-renewal signature. This analysis led us to the unexpected finding that some immune-related genes are in type A (open) compartments. By integrating these findings with RNA-seq datasets for 76 patient-derived primary GSC cultures, we found that *CD276* is expressed at higher levels in GSCs than in their matched bulk GBM samples or in non-neoplastic brain. CD276 was shown to have immune-modulatory functions, but it has not been associated with self-renewal properties before. A previous report showed that the toll-like receptor 4 (TLR4) - which is involved in activating the immune system - is downregulated in GSCs and expressed more highly in their non-GSC counterparts. In non-GSCs, TLR4 repressed a self-renewal network mediated by retinoblastoma binding protein 5 (RBBP5) and ultimately resulted in suppression of the stemness genes *SOX2*, *NANOG* and *OCT4*. These findings suggest that there might be immuno-modulatory proteins like TLR4 that repress self-renewal networks and others like CD276 that positively contribute to stemness. CD276 is expressed in several tumor types and in the tumor vasculature, and it was proposed as a promising therapeutic target (Seaman et al., 2017). Recent preclinical studies showed that an m276-PBD ADC was effective at drastically reducing or even abrogating tumor growth in several cancer models. Here, we show that m276-PBD can be used to target CD276^+^ GSCs to curb their self-renewal potential. This is an exciting result because it suggests that this ADC could be added to current treatments for GBM patients to specifically target and eliminate self-renewing cells. Of note, m276-PBD kills cells in a cell cycle-independent manner. This is important, because self-renewing cells *in vivo* have been shown to be relatively quiescent (Lan et al., 2017), partially explaining their resistance to radiotherapy and the alkylating agent temozolomide that are currently used to treat GBM patients. We propose that adding m276-PBD to the current standard of care might result in the eradication of self-renewing cells and slow recurrences. Beside for m276-PBD, other strategies have been developed to target CD276^+^ cells in cancer. One of them, a CD3-CD276 bimodal antibody, is currently in clinical trials for several solid malignancies, including neuroblastoma and head and neck cancer. Adding GBM patients to these clinical trials could be beneficial for patients. Additionally, chimeric antigen receptor (CAR) T cells could be used to target CD276, exploiting the several-fold higher expression of CD276 in GSCs compared to non-neoplastic brain. The identification of CD276 as a putative new therapeutic target in GBM provides proof-of-principle that integration of structural genomics and other genomic datasets can be successfully integrated to delineate self-renewal states and their vulnerabilities.

## ACKNOWLEDGMENTS

Funding for this work was provided by the Canadian Institutes of Health Research (CIHR) early career award (Institute of Cancer Research) to MG (ICT-156651); a Cancer Research Society Scholarship for the Next Generation of Scientists grant to MG; a Clark Smith Scholarship to MJJ. Research supported by SU2C Canada Cancer Stem Cell Dream Team Research Funding (SU2C-AACR-DT-19-15) provided by the Government of Canada through Genome Canada and the Canadian Institute of Health Research, with supplemental support from the Ontario Institute for Cancer Research, through funding provided by the Government of Ontario. Stand Up To Cancer Canada is a Canadian Registered Charity (Reg. # 80550 6730 RR0001).

